# Observation of the protein-inorganic interface of ferritin by cryo-electron microscopy

**DOI:** 10.1101/2024.09.04.611303

**Authors:** Sagnik Sen, Amar Thaker, Dewight Williams, Po-Lin Chiu, Brent L. Nannenga

## Abstract

Visualizing the structure of the protein-inorganic interface is critically important for our more complete understanding of biomineralization. Unfortunately, there are limited approaches for the direct and detailed study of biomolecules that interact with inorganic materials. Here we use single particle cryo-EM to study the protein-nanoparticle interactions of human light chain ferritin and visualize the high-resolution details of the protein-inorganic interface. In this work, we determined the 2.85 Å structure of human light chain ferritin bound to its native iron oxide nanoparticle substrate. The resulting cryo-EM maps confirmed and enhanced previously proposed interactions of the protein with the material along the B-helix, and revealed new interaction at the C-terminus of light chain ferritin. This work sheds new light on the mechanisms of ferritin biomineralization and further demonstrates the application of cryo-EM for the study of protein-inorganic systems.

## Introduction

The interaction of biomolecules with inorganic materials is an essential process in living systems and has wide ranging impacts in nature, human health, and nanobiotechnology. To understand the fundamental mechanisms of protein-inorganic interactions, and to design and engineer novel hybrid biological-inorganic systems, detailed studies of the protein-inorganic interface are of paramount importance. The proteins responsible for controlling biomineralization and binding to material surfaces often contain regions of high charge; however, the structural organization of the protein at the inorganic interface is challenging to elucidate, due in large part to the difficulties of using traditional structural methods on these protein-inorganic systems^1-11^. As a new approach, the application of single particle cryo-electron microscopy (cryo-EM) to the study of protein-inorganic nanoparticle complexes has attracted attention, and in previous work, cryo-EM has been used to determine the structures of inorganic nanoparticle bound protein complexes^12-15^.

Ferritin is one of the most well-studied biomineralization proteins^16^, and also serves as an excellent model for single particle cryo-EM where it produces very high-resolution reconstructions^17^. Ferritins are ubiquitous proteins that play key roles in iron homeostasis in organisms^18-20^. In addition to their role in nature, ferritin has attracted significant attention for use in various other applications including nanomaterial synthesis and drug delivery^21-24^. Mammalian ferritins are 24-mers composed of both heavy and light chains, and both heavy and light chain ferritins are composed of a four helical bundle (helices A-D) with a fifth short helix (helix E) at the C-terminus that forms the 4-fold axes of the cage. The heavy chain of ferritin has ferroxidase activity for the conversion of Fe^2+^ to Fe^3+^, and the light chain contains a nucleation site for the iron oxide mineral^25, 26^. In crystallographic studies on the interactions of light chain ferritin with iron, it has been shown that indeed the nucleation site consists of residues E60, E61, and E64, and during early events in biomineralization, these form a tri-iron cluster with oxygen atoms^25^. Residue E57 also interacts with iron atoms within this cluster^25, 27^, and a subsequent study suggests E140 may be involved with mineral growth following nucleation^27^. In cryo-EM studies of an iron oxide binding peptide-light chain ferritin fusion, residue H53 was also found to interact with a ferritin synthesized iron oxide nanoparticle (NP). However, the presence of the mineralization peptide shifted the location of the NP away from the nucleation site, making conclusions on the wild-type light chain ferritin interface in this region difficult. While these previous results provide important details on the early events in iron oxide NP nucleation within light chain ferritin, the details of NP growth and native interactions of the protein with the synthesized iron oxide NP have thus far remained elusive.

In this work, we use single particle cryo-EM to study the protein-inorganic interface of human light chain ferritin (HuLF) iron oxide nanoparticle complex. Through the 2.85 Å reconstruction of the NP bound HuLF, the protein-mineral interface along helix B was resolved, and new interactions of the HuLF C-terminus were uncovered. This further demonstrates that cryo-EM can indeed be used to directly visualize the protein-inorganic interface at high-resolution.

## Results and Discussion

To investigate the structure of iron oxide NP bound HuLF, with the ultimate goal of directly observing the protein-inorganic interface, we first optimized the HuLF mineralization process for single particle cryo-EM analysis. Ferritin purified from native sources and ferritin loaded with high amounts of iron in vitro contains large polycrystalline iron oxide cores that are not suitable for high-resolution averaging and 3D reconstruction, as the core is very large and heterogenous^28, 29^. In contrast, the ideal iron oxide nanoparticle HuLF complexes (HuLF-NP) for single particle analysis would have a single nanoparticle within each HuLF cage, representing early-stage nanoparticle nucleation and growth of the first NP within the HuLF cage. To investigate the effects of mineralization conditions on iron oxide NP size and number, we incubated demineralized recombinantly expressed HuLF at increasing ratios of Fe^2+^ to HuLF monomer (1:1, 3:1, and 10:1). Following the reaction, the sample was repurified using size exclusion chromatography (SEC), and small sets of cryo-EM images of the resulting mineralization reactions were collected (Fig. S1). As the ratio of Fe^2+^ was increased, the average size of the resulting NPs increased slightly (2.45 ± 0.44 nm, 2.55 ± 0.45 nm, and 2.63 ± 0.63 nm, for ratios of 1:1, 3:1, and 10:1, respectively), while the number of iron oxide nanoparticles per HuLF increased considerably (Fig. 1A). This suggests that as the nanoparticles reach a certain size, the nucleation of additional nanoparticles is more favorable relative to continued nanoparticle growth. This is consistent with the understanding of ferritin mineralization where multiple nanoparticles are nucleated on the inner surface of the protein cage and grow together to form the final fully loaded ferritin iron complex^28, 29^.

**Fig. 1.**
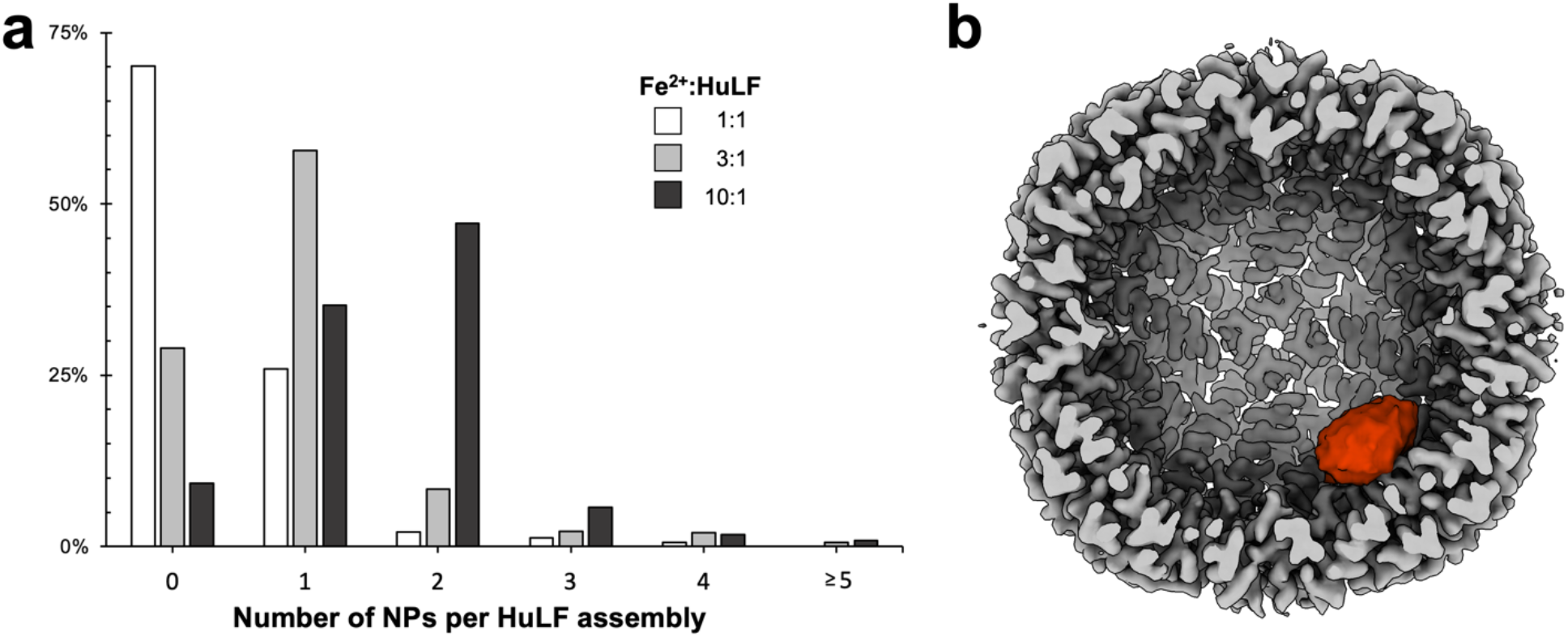
Synthesis and structure of iron oxide NP bound HuLF. (a) The synthesis of NP bound HuLF was optimized to produce the highest ratio of HuLF assemblies with a single NP. As Fe^2+^ was increased, the number of NPs per HuLF were increased with the optimal Fe^2+^:HuLF ratio tested being 3:1. (b) Using the 3:1 ratio, single particle cryo-EM was used to determine the structure of HuLF bound to an iron oxide NP (HuLF-NP) at 2.85 Å, which was sufficient to resolve interactions of the interfacial amino acids.

The 3:1 ratio was selected for subsequent structure determination efforts as it produced the highest level of single nanoparticle HuLF-NP complexes compared with the other ratios tested. High-resolution cryo-EM images were collected and several rounds of 2D classification were employed to curate particles and separate them into groups that contained no nanoparticles (apo-HuLF) or single nanoparticle HuLF-NP complexes (Fig. S2). The classes with poor quality particles or with multiple nanoparticles were removed from subsequent image processing. This process produced a set of 123,544 HuLF particles with single iron oxide NPs, which were used for subsequent 3D reconstruction. Because the presence of the iron oxide NP breaks the octahedral symmetry of the HuLF, reconstructions were performed using C1 symmetry. Additionally, due to the size variation and nonlinearity of the iron oxide NP cryo-EM density, it was necessary to mask out the NP in the final stages of refinement to achieve a high-resolution reconstruction. The results of this image processing workflow (Fig. S2, Fig. S4, and Table S1) ultimately produced a 2.85 Å reconstruction of HuLF-NP (Fig. 1B), with the local resolution highest in the core of the helical HuLF structure and lower in the outer areas of the NP (Fig. S5). Additionally, a 2.10 Å structure of apo-HuLF was determined using standard octahedral symmetry, which provided a comparison with the HuLF-NP structure.

The high-resolution reconstruction of the HuLF-NP complex provides a direct observation of the interface between the protein and the inorganic NP, revealing the details of the amino acids responsible for the interactions between HuLF and the inorganic material (Fig. 2). The major protein-inorganic interface is formed by helix B of HuLF. The iron oxide NP is positioned directly over helix B and residues E60, E61, and E64 (Fig. 2), which have been shown to provide the initial scaffold for the nucleation of the iron oxide material^25^. The cryo-EM map shows that these residues maintain their interaction with the formed NP, further demonstrating that this is indeed the site of iron oxide nucleation, as well as the site for NP growth and binding on the inner surface of the HuLF cage. It is important to note that the density for these residues is poor in the other non-NP interacting chains of HuLF-NP (Fig. S6a-b), as well as in the apo-HuLF structure determined from the same sample (Fig. S6c-d). The lack of density for these residues is also seen in high-resolution cryo-EM reconstructions of apo-HuLF and horse light chain apoferritin from other studies^30, 31^. However, in the presence of the NP, the density for these residues is clear in the complex demonstrating that the positions of the amino acids are stabilized, and the signal is improved due to their interactions with the surface of the iron oxide NP. In addition to E60, E61 and E64, E57 can also be seen interfacing with the iron oxide NP. Through crystallographic studies of early iron oxide nucleation, this residue was previously hypothesized to bind and shuttle iron ions to the E60, E61, E64 nucleation site^25, 27^. Here, the cryo-EM map of the HuLF-NP complex shows that E57 is also part of the binding interface with the formed iron oxide NP. As with the other glutamic acid residues on helix B, E57 is also not well resolved in the absence of NPs (Fig S6), however the density becomes clear when bound to the iron oxide material.

**Fig. 2.**
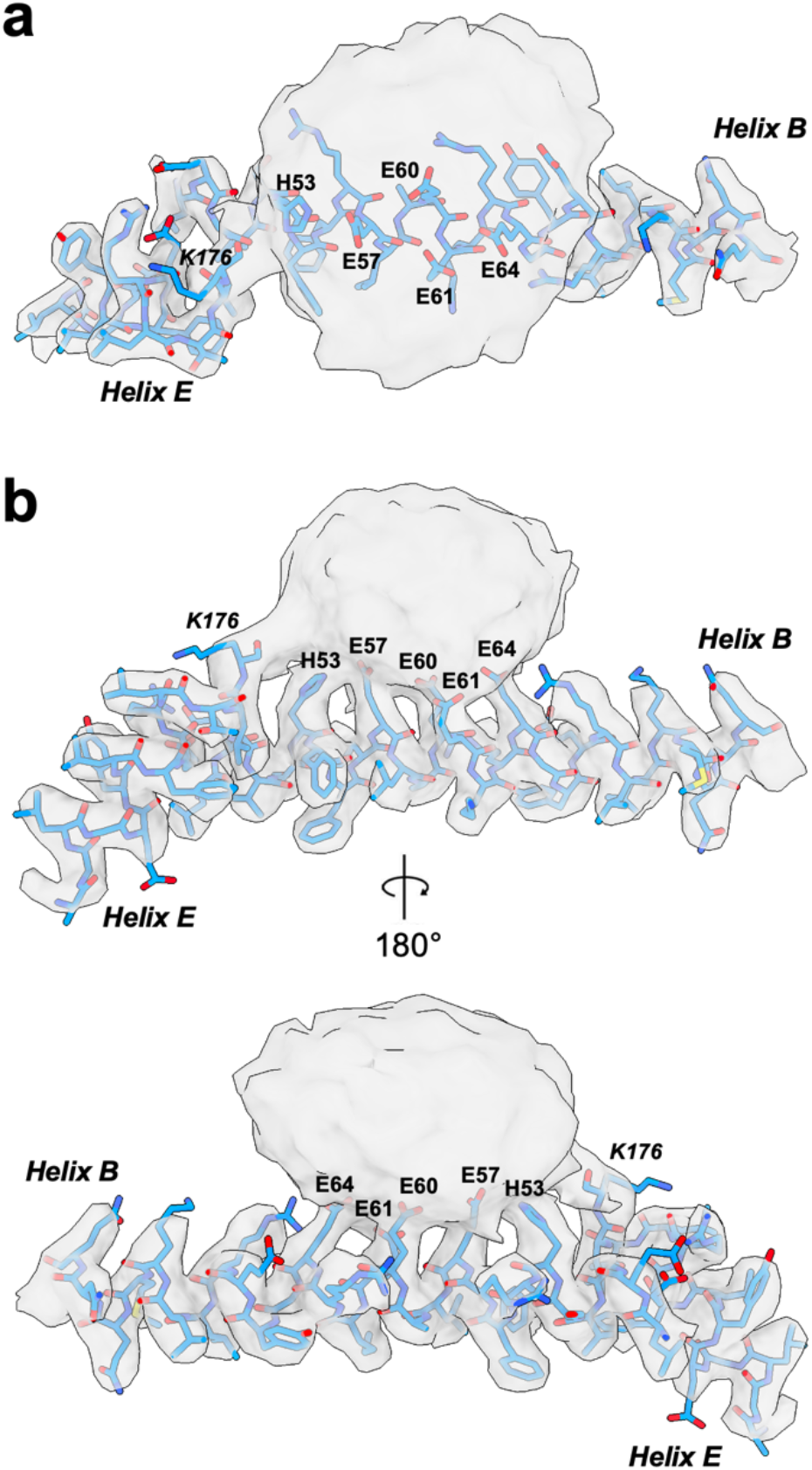
The cryo-EM structure of the protein-inorganic interface of HuLF. The 2.85Å cryo-EM map and model of the HuLF-NP complex is shown from the top (a) and sides (b). The cryo-EM map is contoured at 5 σ around helix B and E of the HuLF-NP model, which is where the protein-inorganic interface is formed. The key residues in the interaction with helix B are labeled (H53, E57, E60, E61, and E64) as well as the interaction of the C-terminus, which follows helix E and is labeled with the last residue modeled, K176.

While the cryo-EM maps of the HuLF-NP complex confirms much of what was previously found by the X-ray crystallographic studies on HuLF and initial iron clusters^25,27^, there are key differences seen between the structures on helix B (Fig. 3). The structures of helix B between the two approaches are very similar, with an overall RMSD of 0.405 Å for helix B from the cryo-EM model presented here and the previous X-ray structure of HuLF bound to a tri-iron cluster (PDB ID: 5lg8)^25^. When the two structures are aligned, the relative distances between the Cα atoms of E57, E60, E61, and E64 are 0.37 Å, 0.14 Å, 0.22 Å, and 0.29 Å, respectively (Table S2). However, for residues E57 and E60, the overall orientations of the side chains bound to the nanoparticle in cryo-EM model are different relative to the tri-iron cluster structure determined by X-ray crystallography (Fig. 3), with the relative Cδ distances of 2.66 Å for E57 and 1.72 Å for E60 (Table S2). When examining the cryo-EM maps near these residues, it is clear the orientations of E57 and E60 from the tri-iron bound X-ray structure do not match the HuLF-NP complex structure (Fig. 3C). The difference in the position of E61 is less drastic, with the Cγ and Cδ being shifted by 1.29 Å and 1.10 Å, respectively, while the relative orientation of the residue in both structures is similar. E64 is relatively unchanged between the two structures with the differences between each side chain carbon being less than the overall RMSD of helix B. The differences in the side chain positions of the glutamic acid residues, particularly E57, reveals that there is a conformational change of the interface as the growth of the nanoparticle proceeds from the initial iron oxide cluster to the larger nanoparticles in this study. Indeed, the orientation of E57 in the HuLF-NP model is similar to the X-ray structure of the HuLF E60A/E61A/E64A triple mutant that is not capable of forming the tri-iron cluster, but forms an alternative iron cluster between H53 and E57 at a reduced rate^27^. Along with the glutamic acid interactions, H53 on helix B is also shown to interact with the surface of the NP, which is also seen in the alternative iron cluster from the HuLF triple mutant described above^27^. Altogether, the cryo-EM map of HuLF-NP around this group of charged residues, H53, E57, E60, E61, and E64, provides a more complete view of the interface between helix B of HuLF and the more mature iron oxide NP.

**Fig. 3.**
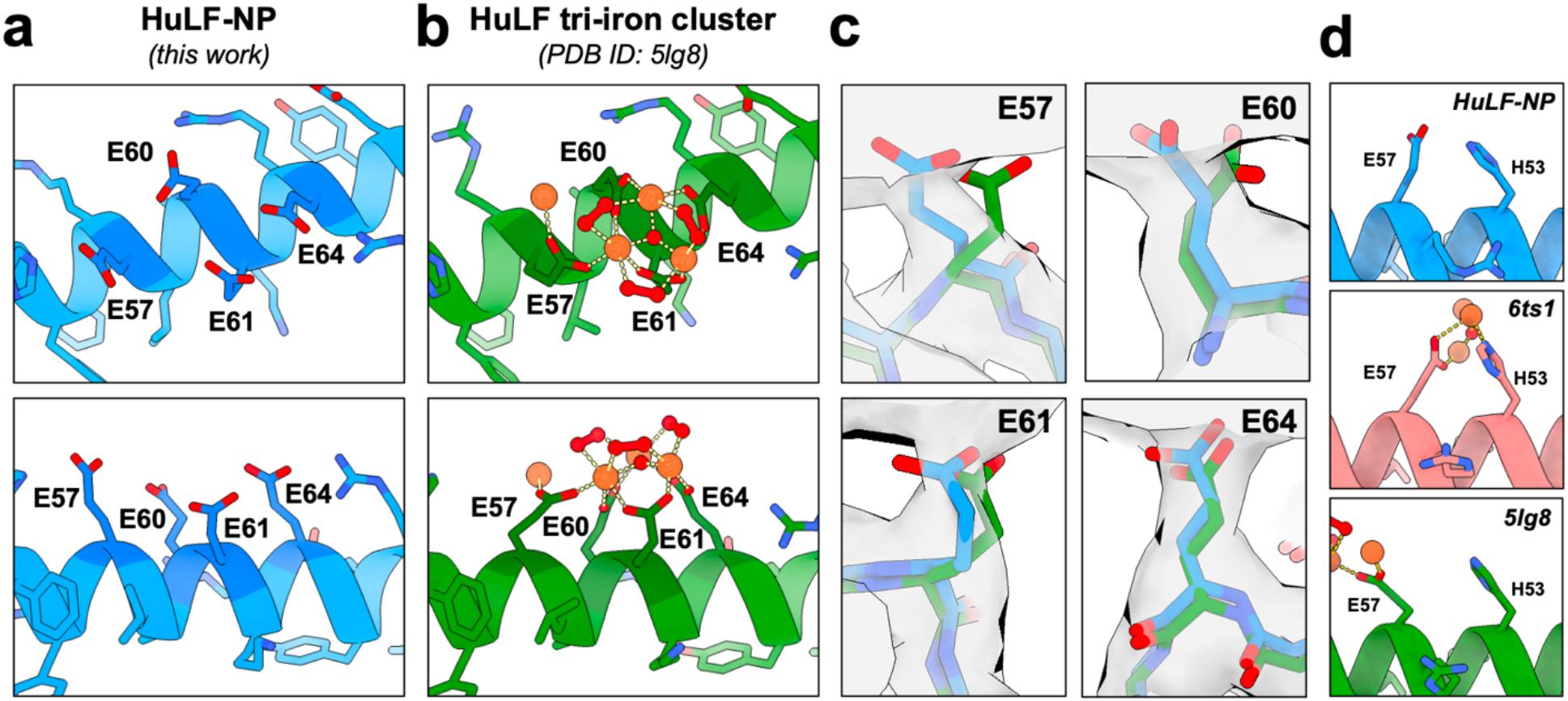
Comparisons of the cryo-EM HuLF-NP complex structure and iron cluster structures determined by X-ray crystallography. The models derived from the cryo-EM maps of HuLF-NP determined in this work (a) show key differences when compared with the model of smaller tri-iron clusters by X-ray crystallography (b, PDB ID: 5lg8), particularly the conformations of E57 and E60 (c). (d) While there are significant differences between E57 in HuLF-NP (d top) and the tri-iron cluster from PDB ID 5lg8 (d bottom), the conformation of E57 in the HuLF-NP structure is much more similar to E57 from the E60A/E61A/E64A HuLF mutant that forms an alternative iron cluster (d bottom. PDB ID: 6ts1). The HuLF-NP maps presented in (c) are contoured at 5 σ.

In addition to the interface formed by helix B, the HuLF-NP complex map sheds light on other interactions of the inner surface of light chain ferritin. As mentioned above, residue E140 was previously shown using X-ray crystallography to be involved in interacting with a larger octa-iron cluster^27^. This interaction was not seen in earlier structures of the smaller tri-iron cluster^25^, which suggested that E140 was part of the interface of the growing iron oxide NP. However, in the HuLF-NP cryo-EM structure presented here, no clear interactions between E140 with the nanoparticle could be seen, showing it is not part of the NP interface and suggesting that this may be a transient interaction during early NP growth. Perhaps the most interesting finding of the HuLF-NP structure was that the cryo-EM map shows further interactions of the HuLF C-terminus with the iron oxide NP (Fig. 2). In other structures of both heavy and light chain ferritin, the C-terminal residues were not well-ordered and were not resolved by X-ray crystallography or cryo-EM^25, 27, 30, 31^. This can also be seen here in the apo-HuLF structure, as well as in the other HuLF chains of the HuLF-NP structure that are not adjacent to the NP (Fig. S6). In each of these cases, there is no continuous density for the C-terminal residues, H177 and D178, of HuLF and the model is truncated at residue K176. However, in the presence of the NP, there is continuous density extending from the interacting HuLF chain’s C-terminus to the iron oxide NP, indicating that the C-terminus interacts with the NP surface. The map in the region of the C-terminus is not detailed enough to allow the clear modeling of H177 and D178 and different potential conformations of these terminal residues could be proposed.

To confirm that the density extending to the iron oxide NP is indeed the terminal H177 and D178 residues of HuLF that are not resolved in the apo structures, a mutant HuLF construct was made which lacks these final two residues (HuLF_Δ177-178_). Iron oxide mineralization and cryo-EM structure determination was performed with the truncated construct using the same approaches as for the wild-type HuLF (Fig. S3, Fig. S4, and Table S1). The resulting 3.3 Å reconstruction of the HuLF_Δ177-178_ NP complex shows no clear density between the iron oxide NP and the HuLF in the region of the truncated C-terminus (Fig. 4), and the map from the apo-HuLF_Δ177-178_ shows a similar reduction in density for the C-terminus (Fig. S7). The maps derived from the truncated construct confirms that the C-terminus of wild-type HuLF does indeed interact with the iron oxide NP. When HuLF is in its unbound state, the C-terminus is unresolved in the cryo-EM maps, implying its flexible and disordered nature. However, in the presence of the iron oxide NP, clear density is seen interfacing with the material. The ordering of the C-terminus in the presence of the NP is consistent with the hypothesis that the process of biomineralization can convert poorly ordered regions of proteins to structured domains through their interactions with inorganic materials^15, 32, 33^.

**Fig. 4.**
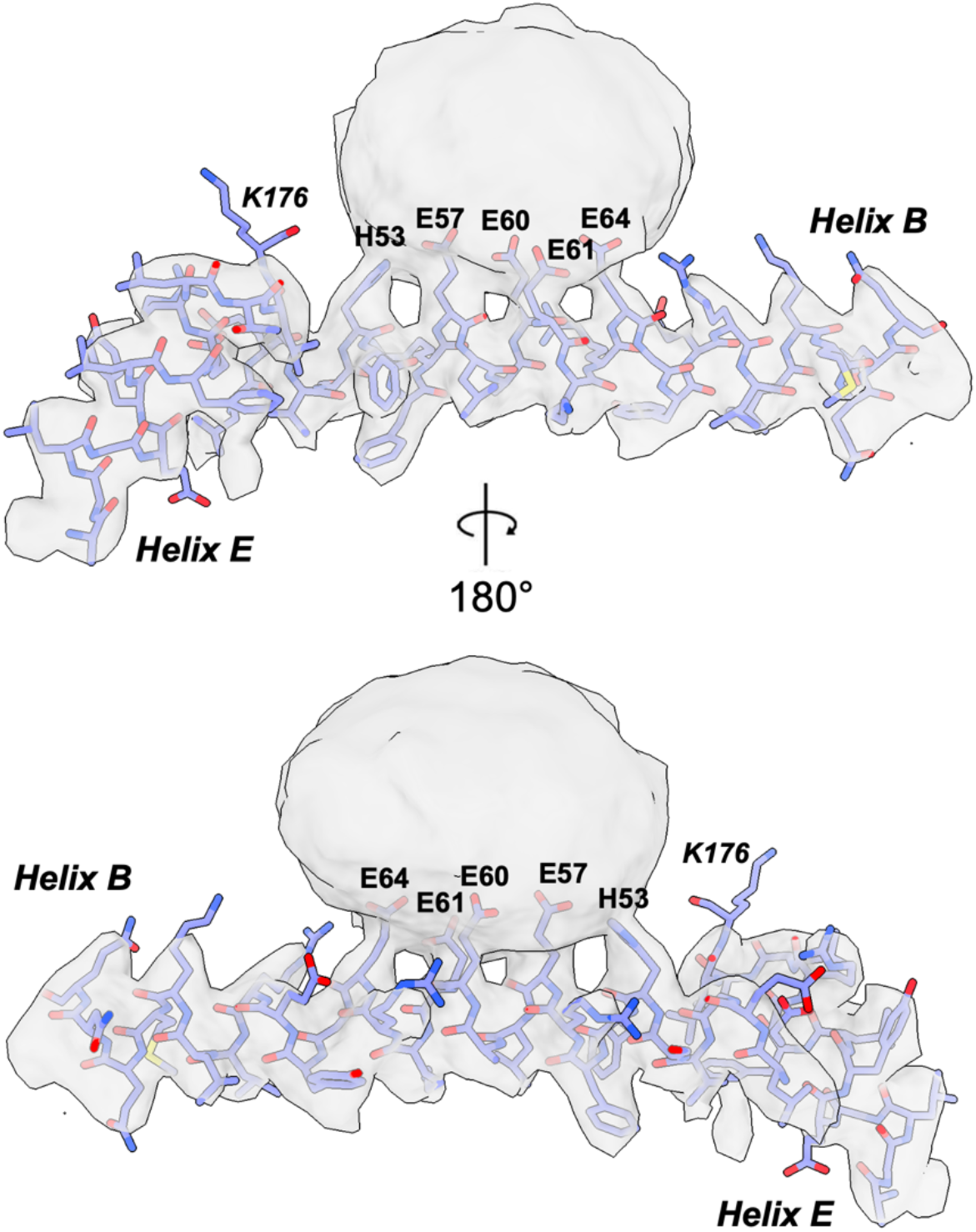
The cryo-EM structure of the NP bound HuLF C-terminal truncated variant. The HuLF_Δ177-178_ construct was used to produce a 3.30 Å cryo-EM map of the NP complex with HuLF lacking the final two residues, H177 and D178. The map viewed from the top (a) and sides (b), is similar to the wild type HuLF-NP structure, with the exception of the lack of continuous density from the C-terminus of HuLF_Δ177-178_ to the NP. The cryo-EM map is contoured at 3.0 σ around helix B and E of the chain A.

Because of the cryo-EM density attributed to the C-terminus, we hypothesized that the removal of the C-terminus would influence iron oxide nanoparticle formation. Measurements of NP size in the cryo-EM micrographs yielded an average longest diameter of 3.05 ± 0.69 nm for NPs synthesized within HuLF_Δ177-178_, which was significantly larger than the measurement of 2.59 ± 0.69 nm for the NPs from the wild-type HuLF (n = 125; *p* = 3.09 × 10^−7^). Additionally, when the oxidation of iron following incubation with Fe^2+^ was measured using absorbance at 350 nm^25^, the values for HuLF_Δ177-178_ were increased relative to the same measurements for wild-type HuLF (Fig. S8). Past studies on the C-terminus of ferritin have focused on the E-helix at the four-fold axis, with deletions that were much larger than the one in this study, which removed the majority of the E-helix^34, 35^. These larger truncations ultimately lead to a reduction of ferritin stability and subsequently a reduced capacity for mineralization. The findings in this work suggest that the C-terminus may have a more important role in biomineralization beyond only enhancing the stability of the overall ferritin structure.

## Conclusion

The cryo-EM structures of iron oxide NP bound HuLF presented here provide a more complete view of the protein-inorganic interface in ferritin, complimenting past studies and adding to our understanding of ferritin biomineralization. Additionally, this work further demonstrates that single particle cryo-EM is capable of resolving high-resolution details at the protein-inorganic interface. Therefore cryo-EM promises to be a unique approach for enhancing our understanding of the structure and function of macromolecules involved in biomineralization.

## Methods

### Ferritin Purification

Human light chain ferritin (HuLF) was cloned into pET-24a(+) using Gibson Assembly. The resulting plasmid was used for protein expression in *E. coli* BL21(DE3). Protein expression was performed using 1 L LB media supplemented with 50 μg/mL kanamycin, 0.5 % glycerol, 0.05% glucose, and 1mM IPTG was used to induce expression at 25°C for 24 hours. Cells were harvested by centrifugation at 8,000 g for 10 mins, and the resulting cell pellets were stored at -20°C.

For HuLF purification, the cell pellets were first thawed in a lysis buffer (150 mM NaCl, 20 mM Tris-HCl (pH 7.3)) with 1 mg/mL Lysozyme and 1mM PMSF added. Lysis was performed using sonication and cell debris was removed by centrifugation. The supernatant was treated with DNAse1 with 50 mM MgCl2 as cofactor at room temperature for 30 minutes. Ferritin being a heat-resistant protein, heat treatment was done on the sample at 75°C for 20 minutes in a heat bath and the denatured proteins were removed by centrifugation. Then the supernatant was subjected to ammonium sulfate (35% saturation) at 4°C for 1 hour. The sample was centrifuged at 22,000 g for 30 minutes at 4°C and the precipitated proteins were removed by collection of the supernatant. The supernatant was then subjected to further ammonium sulfate treatment to a final saturation of 65% and the previous centrifugation step was repeated. The precipitated fraction following centrifugation was recovered. The precipitated fraction containing HuLF was resuspended in a SEC buffer (50 mM NaCl, 20mM Tris-HCl (pH 7.3)). Any bound iron was removed through 4 steps of dialysis with a buffer consisting of 200 mM NaCl, 3 mM EDTA, ammonium thioglycolate (1:400), and 20 mM Tris HCl (pH 7.0). Final SEC purification was performed with a Superose 6 Increase 10/300 GL column. The presence of HuLF was confirmed using SDS-PAGE analysis of the fractions obtained after SEC.

The truncated HuLF_Δ177-178_ was constructed by site-directed mutagenesis of HuLF and was expressed and purified as described for HuLF.

### Iron Nanoparticle synthesis and sample preparation

Iron oxide nanoparticle (NP) formation was performed by adding Mohr’s salt (ammonium iron(II) sulfate) to purified HuLF (20 μM) at a ratio of 3 Fe^2+^ to 1 HuLF monomer, and the reaction was carried out at room temperature for 4 hours. The sample then was centrifuged at 12,000 g for one minute and any precipitated iron oxide and ferritin were removed. The supernatant was passed through a Superdex 200 10/300 GL column and the iron loaded HuLF (HuLF-NP) peak was collected for cryo-EM sample preparation.

For cryo-EM sample preparation, 3 μL of HuLF-NP sample was added to Quantifoil R2/1 grids, blotted for 3 seconds, and plunge-frozen in liquid ethane using a Vitrobot (Thermo Fisher). Grids were imaged in a Titan Krios (300 KeV) cryo-transmission electron microscope (Thermo Fisher) equipped with a K2 Summit direct electron detector (Gatan). Data sets of 5,620 and 2,862 movies were collected in counting mode with a pixel size of 1.03 Å for HuLF and HuLF_Δ177-178_, respectively.

### Cryo EM image processing

Data processing was performed in the cryo-EM software platform CryoSPARC (version 3.3.2)^36^. Movies were imported into cryoSPARC, followed by motion correction and CTF correction.

For HuLF, 5,484 micrographs were selected for further processing after manual curation. Template based particle picking was used to select 2,310,211 particles, which were extracted from the micrographs with a box size of 256 pixels and the ‘Fourier crop to box size’ parameter set at 128 pixels. The Fourier-cropped images were used for 2D classification and curation. For the HuLF nanoparticle complex, all 2D classes showing a single nanoparticle were selected and used to calculate a single *ab initio* 3D volume. This was refined using no symmetry (C1 symmetry) and the resultant volume was used to re-extract the particles using the ‘recenter using aligned shifts option’. The particle stack was then subjected to 6 cycles of 2D classification to produce a final stack of 123,544 particles. This was used to obtain a single *ab initio* 3D class at moderate resolution. Using UCSF Chimera (version 1.16), the isosurface where only the nanoparticle density was visible was used to create a binary mask around the nanoparticle using the ‘vop’ tool. This mask was imported back into CryoSPARC and the ‘Particle subtraction’ job was used to subtract the density of the nanoparticle from the individual particle projections. Refinement was then performed with a reference of the 25 Å map of HuLF. The resultant volume containing the HuLF-NP complex was further refined with local and global CTF parameters and the final 2.85 Å map was obtained.

The map of apo-HuLF from the same sample was determined using the 2D classes that did not show the presence of an iron oxide NP. The apo classes were initially put through 6 rounds of 2D classification and the resultant stack was used to obtain a single 3D ab initio volume using octahedral ‘*O*’ symmetry. The particles were re-extracted using the poses previously aligned and were put through another 5 rounds of 2D classification. The curated particles were used to generate a single 3D reconstruction. This was again refined using *O* symmetry and after further corrections for local and global CTF parameters the final 2.10 Å apo-HuLF map was obtained.

Images of the truncated HuLF_Δ177-178_ construct were processed in the same manner as described for the wild type HuLF samples above. 2,862 micrographs were selected, and 746,972 particles were initially extracted. For NP bound HuLF_Δ177-178_ a total of 85,039 particles were used in the final 3.30 Å reconstruction, again using C1 symmetry and nanoparticle subtraction. For the apo-HuLF_Δ177-178_ 381,609 particles were used for the 2.36 Å reconstruction.

Model building was performed using COOT^37^ and ISOLDE^38^ within ChimeraX^39^. The Phenix suite was used for map and model validation and local resolution calculations^40^. Map and model visualization was performed using ChimeraX.

### Nanoparticle and iron oxidation measurements

Motion corrected cryo-EM micrographs were used to analyze the number and size of iron oxide NPs within the HuLF and HuLF_Δ177-178_ using ImageJ^41^. The oxidation of Fe^2+^ to Fe^3+^ was quantified by measuring the absorbance at 350 nm. Identical reactions of HuLF and iron that were used for cryo-EM sample preparation were prepared as described above. The absorbance of the samples following 4 hours of incubation at room temperature were measured using a SpectraMax M5 plate reader (Molecular Devices)

## Data Availability

Maps and models in this study were deposited with the EMDB and PDB under the following accession numbers: PDB ID 9bpj, EMD-44779; PDB ID 9bpk, EMD-44780; PDB ID 9bpi, EMD-44778; and PDB ID 9bq5, EMD-44797; for HuLF-NP, apo-HuLF, HuLF_Δ177-178_ NP, and apo-HuLF_Δ177-178_, respectively.

## Acknowledgements

BLN would like to acknowledge support from the National Science Foundation (DMR-1942084). We would like to acknowledge the use of the Titan Krios at the Eyring Materials Center at Arizona State University and the funding of this instrument by NSF MRI 1531991.

