## Supplemental Information for "Observation of the protein-inorganic interface of ferritin by cryo-electron microscopy"

### Supplementary Data

#### Table S1. Cryo-EM data collection, refinement, and validation statistics

| | **HuLF-NP**<br>PDB ID 9bpj<br>EMD-44777 | **apo-HuLF**<br>PDB ID 9bpk<br>EMD-44780 | **$HuLF_{\Delta 177-178}$ NP**<br>PDB ID 9bpi<br>EMD-44778 | **apo-$HuLF_{\Delta 177-178}$**<br>PDB ID 9bq5<br>EMD-44797 |
|---|---|---|---|---|
| **Data Collection and Processing** | | | | |
| Magnification | 48,544 | 48,544 | 48,544 | 48,544 |
| Voltage (kV) | 300 | 300 | 300 | 300 |
| Exposure ($e^-/Å^2$) | 72.48 | 72.48 | 72.48 | 72.48 |
| Defocus range (μm) | -0.8 to -2.6 | -0.8 to -2.6 | -0.8 to -2.6 | -0.8 to -2.6 |
| Pixel size (Å) | 1.03 | 1.03 | 1.03 | 1.03 |
| Symmetry | C1 | O | C1 | O |
| Final particle images | 123,544 | 542,755 | 85,039 | 381,609 |
| Map resolution (FSC = 0.143) (Å) | 2.85 | 2.10 | 3.30 | 2.36 |
| **Model Refinement** | | | | |
| Model resolution (Å) | | | | |
| FSC threshold (0.5 / 0.143) | 3.27 / 2.85 | 2.29 / 2.10 | 3.88 / 3.30 | 2.57 / 2.36 |
| Model composition | | | | |
| Non-hydrogen atoms | 33,192 | 33,214 | 33,192 | 33,214 |
| Protein residues | 4,128 | 4,128 | 4,128 | 4,128 |
| RMS deviations | | | | |
| Bond lengths (Å) | 0.011 | 0.011 | 0.011 | 0.011 |
| Bond angles (°) | 1.909 | 1.907 | 1.868 | 1.914 |
| Validation | | | | |
| MolProbity Score | 1.22 | 1.19 | 1.52 | 1.30 |
| Clash Score | 0.30 | 0.65 | 0.50 | 0.96 |
| Ramachandran Plot | | | | |
| Favored (%) | 96.18 | 97.72 | 94.75 | 97.60 |
| Allowed (%) | 3.14 | 2.01 | 4.49 | 2.08 |
| Outliers (%) | 0.69 | 0.27 | 0.76 | 0.32 |

#### Table S2. Relative distances between key glutamic acids in HuLF-NP and previous models

| | HuLF-NP:5lg8 | | | | HuLF-NP:6ts1 |
|---|---|---|---|---|---|
| | E57 | E60 | E61 | E64 | E57 |
| **Cα - Cα (Å)** | 0.37 | 0.14 | 0.22 | 0.29 | 0.47 |
| **Cβ - Cβ (Å)** | 0.67 | 0.13 | 0.21 | 0.30 | 0.22 |
| **Cγ - Cγ (Å)** | 2.51 | 0.30 | 1.29 | 0.36 | 0.41 |
| **Cδ - Cδ (Å)** | 2.66 | 1.72 | 1.10 | 0.36 | 1.11 |

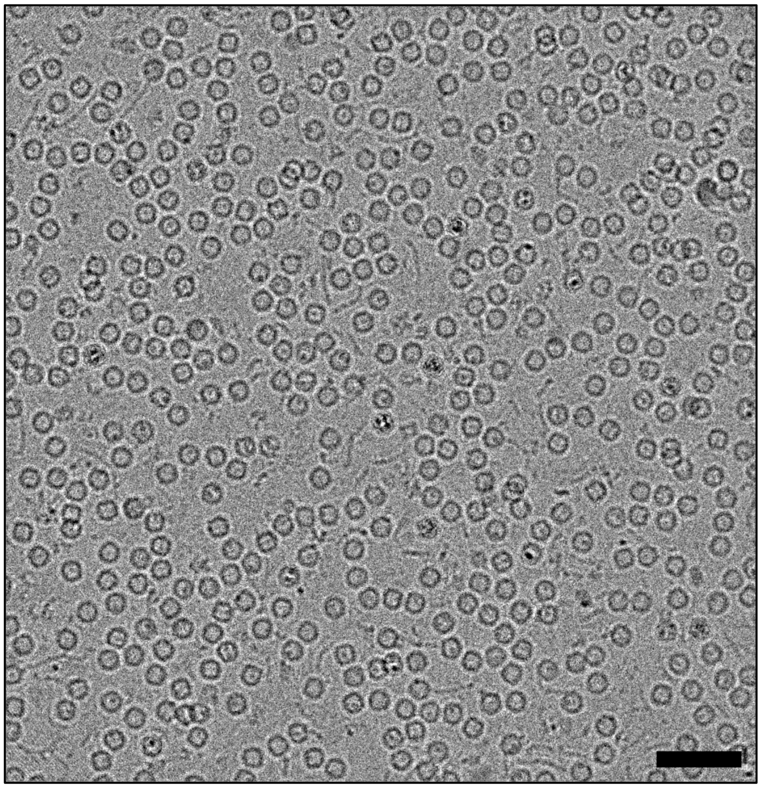

**Fig. S1. Representative cryo-EM micrograph of iron oxide NP bound HuLF.** Scale bar represents 50 nm.

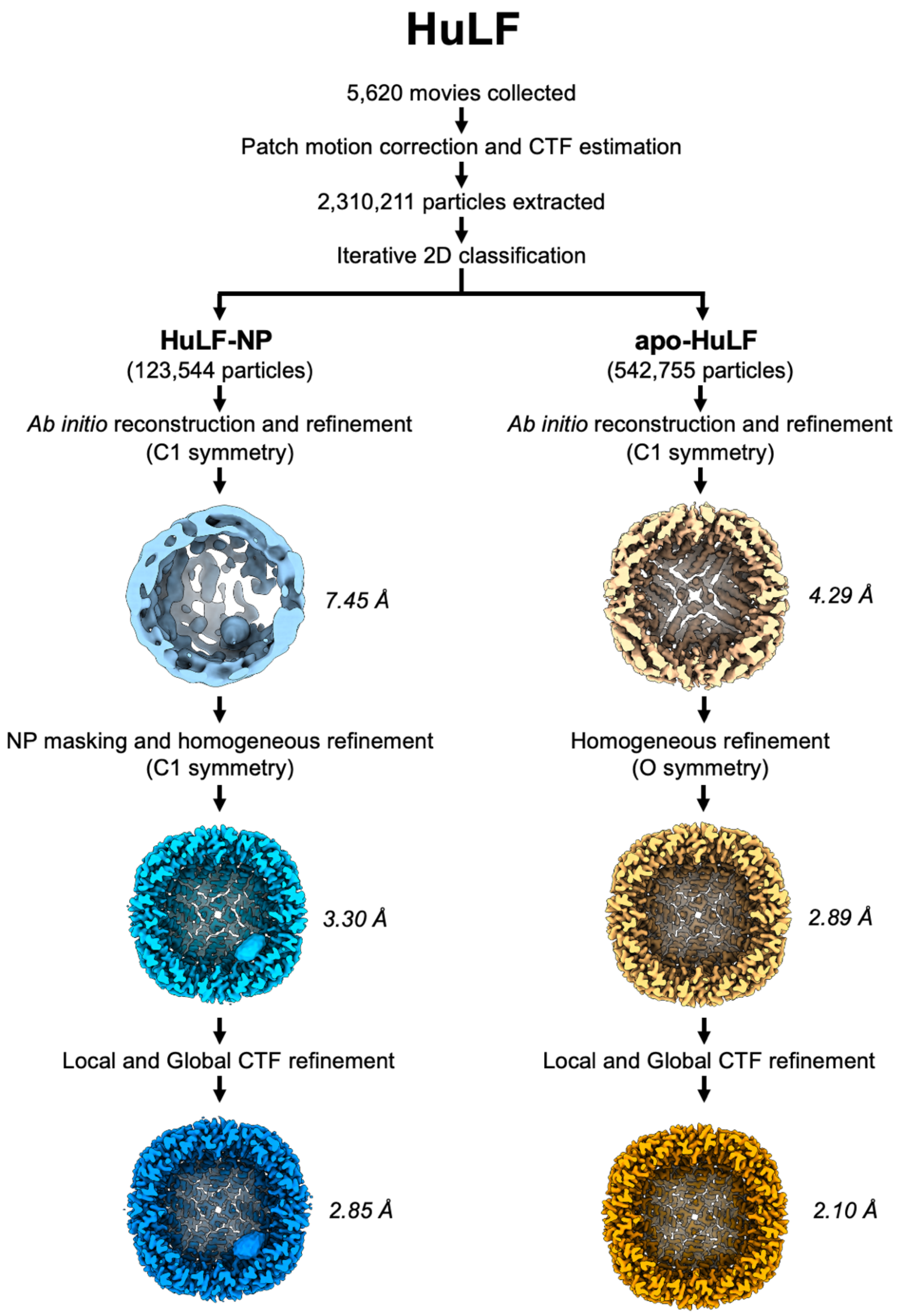


**Fig. S2. Data processing workflow for HuLF**

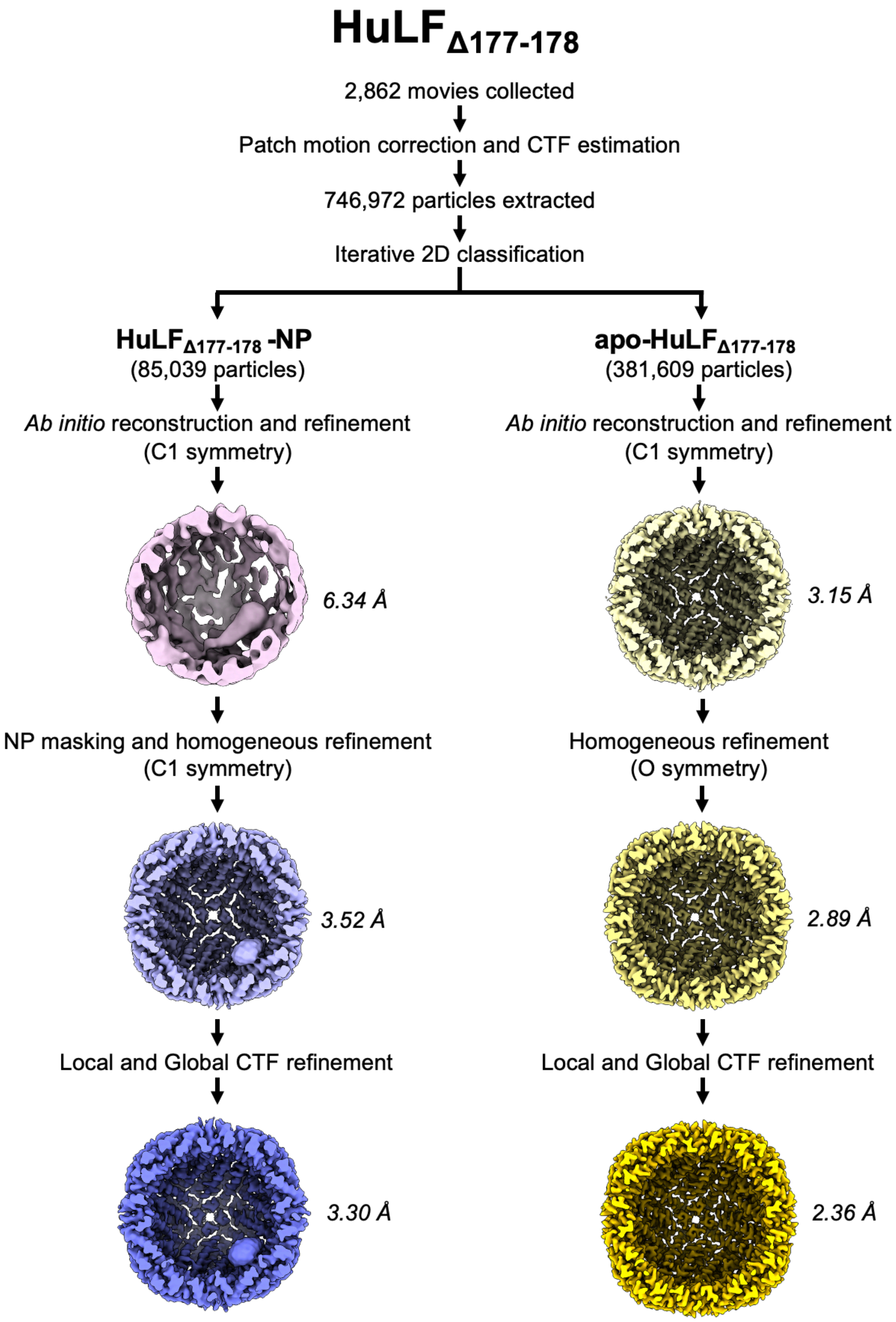


**Fig. S3. Data processing workflow for HuLF$_{\Delta 177\text{-}178}$**

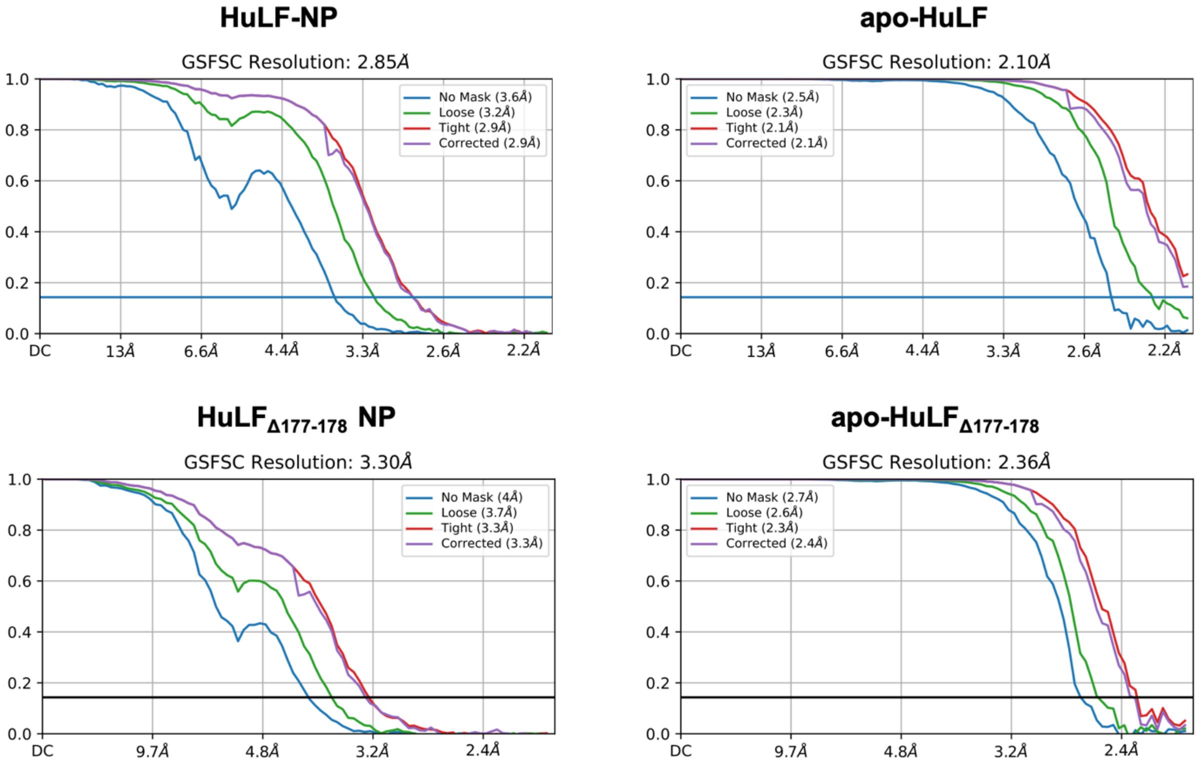


**Fig. S4. Fourier-shell correlation (FSC) plots for each of the cryo-EM reconstructions**

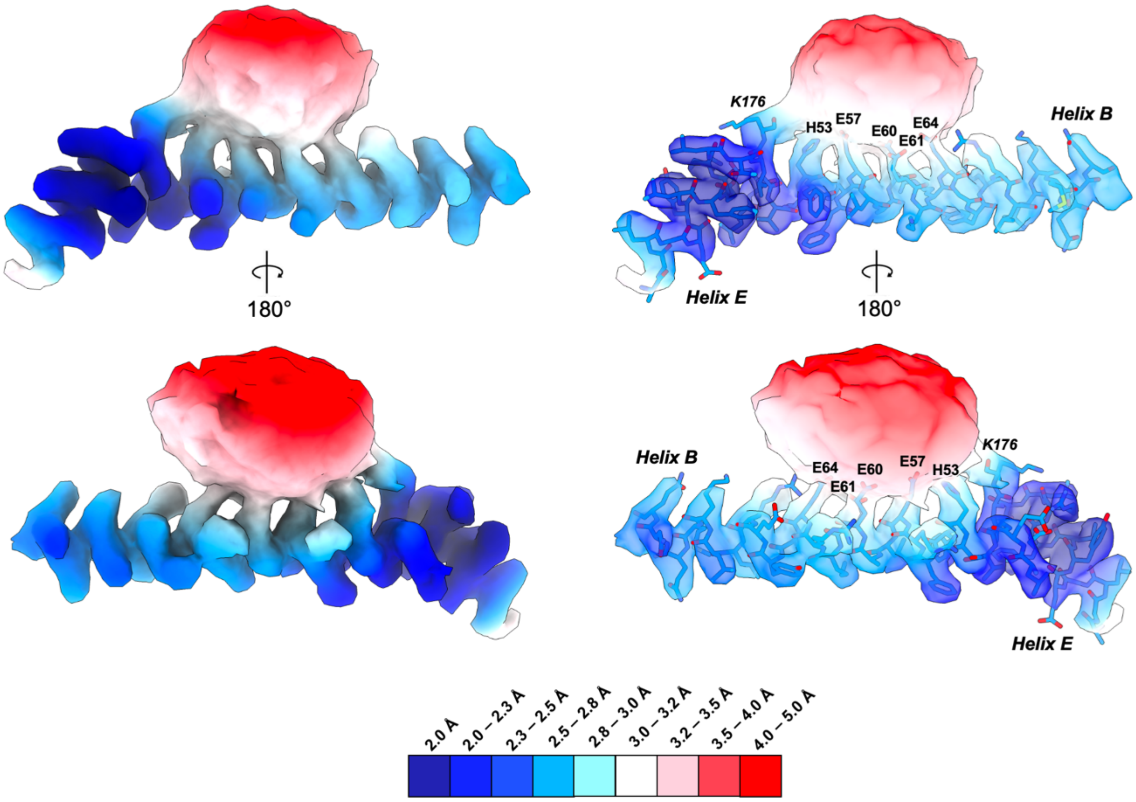


**Fig. S5. Local resolution map around the protein-inorganic interface of HuLF-NP.** Local resolution map was calculated using Phenix and visualized using ChimeraX.

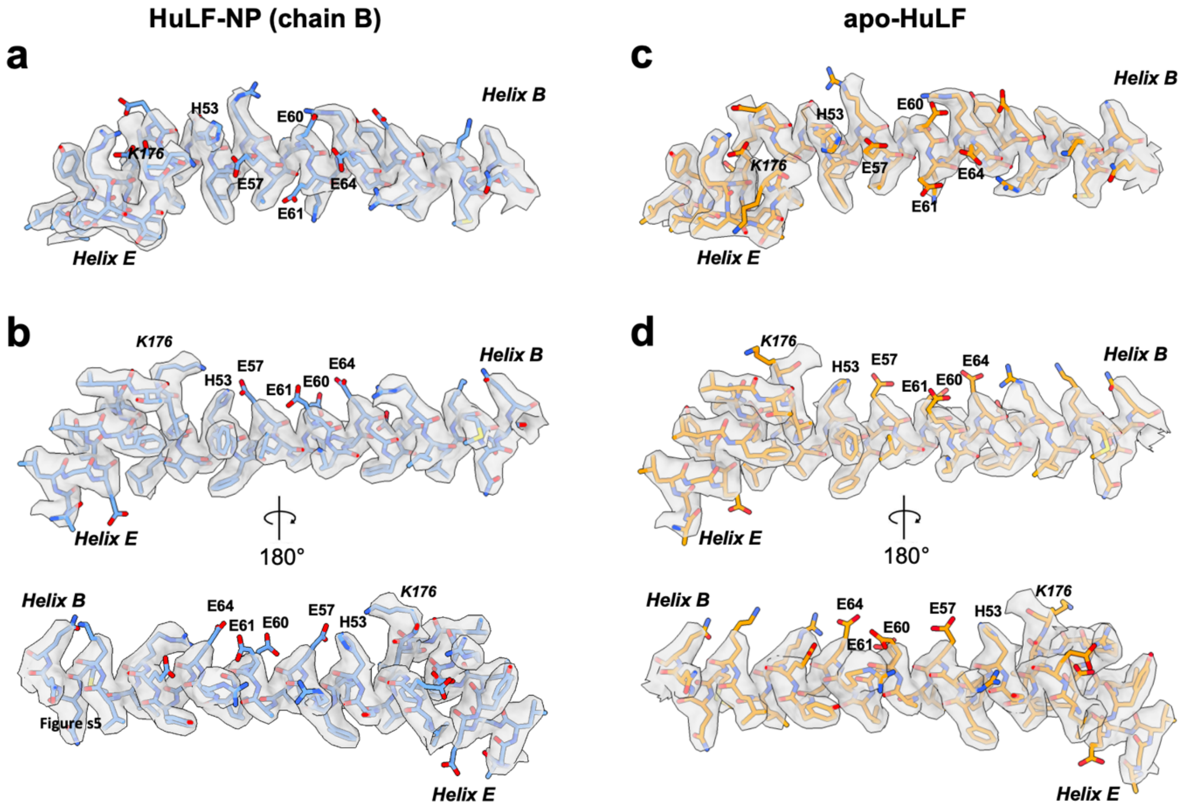


**Fig. S5. Cryo-EM maps of HuLF not interacting with iron oxide NPs.** (a-b) Top and side views of a representative chain (chain B) in the HuLF-NP structure that is not interacting with the NP (chain A). (c-d) Top and side views of apo-HuLF in the area that the NP would be bound (see Fig. 2). The structure of apo-HuLF is from the same data set used to determine HuLF-NP (see Fig. S2). Maps are contoured at 5.0 σ.

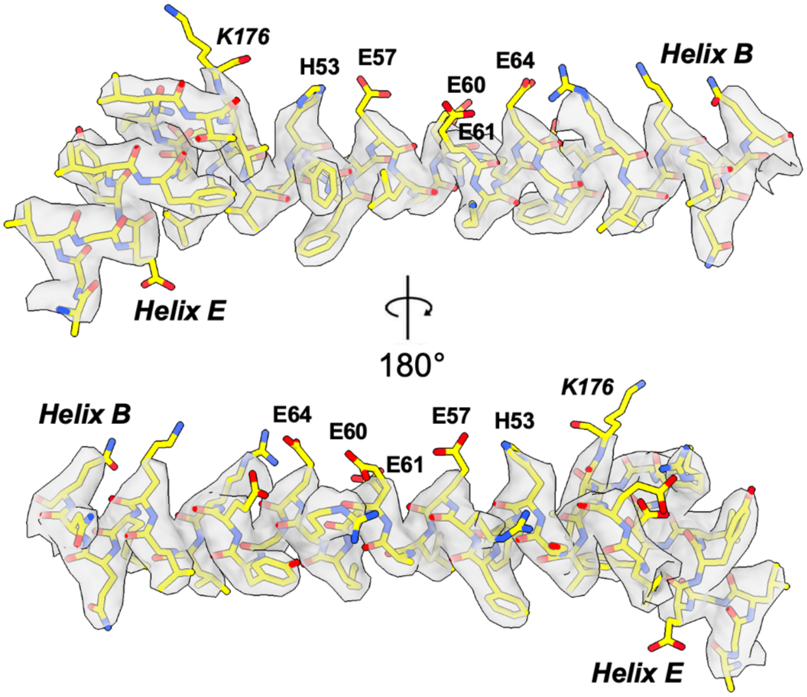


**Fig. S6. Cryo-EM map of apo-HuLF$_{\Delta 177\text{-}178}$.** (a-b) Top and side views of apo-HuLF$_{\Delta 177\text{-}178}$ in the area that the NP would be bound (see Fig. 4). Maps are contoured at 3.0 σ.

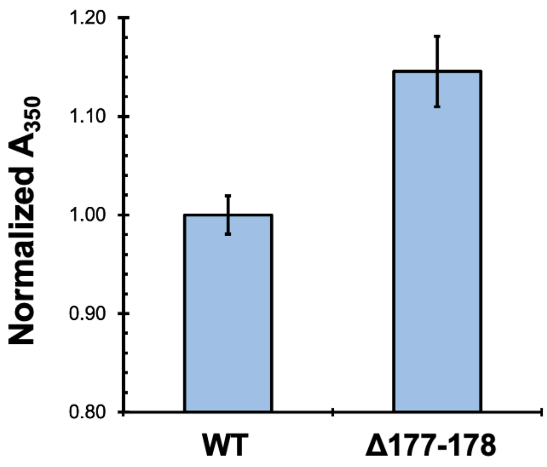


| Average NP size | |
|---|---|
| **WT** | 2.59 ± 0.69 nm |
| **Δ177-178** | 3.05 ± 0.69 nm |

*n = 125; p = 3.09×10⁻⁷*

**Fig. S7. Total Iron oxidation and NP size differences between HuLF and HuLF$_{\Delta 177\text{-}178}$.** The total amount of Fe2+ oxidation to Fe3+ was determined by measuring the absorbance at 350 nm. The reaction in the presence HuLF$_{\Delta 177\text{-}178}$ led to higher A350 relative to the wild type HuLF. When the nanoparticle size was measured using the cryo-EM micrographs, it was found that the nanoparticle size was significantly larger for the truncated variant relative to the wild type HuLF.
